# Human Alveolar and Monocyte-derived Human Macrophage Responses to Mycobacterium tuberculosis

**DOI:** 10.1101/2024.02.20.581265

**Authors:** Monica Campo, Kimberly A. Dill-McFarland, Glenna J. Peterson, Basilin Benson, Shawn J. Skerrett, Thomas R. Hawn

**Affiliations:** Department of Medicine, University of Minnesota, Minneapolis, MN; Department of Medicine, University of Washington, Seattle, WA; Systems Immunology Program, Benaroya Research Institute, Seattle, WA

**Author notes:** Corresponding Author: Monica Campo, MD. Author Contributions: M.C., T.R.H., designed the studies. G.J.P. performed the experiments. K.A.D. and B.B. performed the RNA-sequencing analysis. S.J.S. provided alveolar macrophages. M.C. and T.R.H. wrote the manuscript. M.C., K.A.D., G.J.P., B.B., S.J.S., and T.R.H., edited and revised the manuscript.

**Keywords:** Human alveolar macrophages, monocyte derived macrophages, *Mycobacterium tuberculosis*, interferon, IFNA8

## Abstract

**Rationale:** Alveolar macrophages (AMs) and recruited monocyte-derived macrophages (MDMs) mediate early lung immune responses to *Mycobacterium tuberculosis* (Mtb). Differences in the response of these distinct cell types is poorly understood and may provide insight into mechanisms of TB pathogenesis.

**Objectives:** To determine whether Mtb induces unique and essential anti-microbial pathways in human AMs compared to MDMs.

**Methods:** Using paired human AMs and 5-day MCSF-derived MDMs from 6 healthy volunteers, we infected cells with Mtb H37Rv for 6 hours, isolated RNA, and analyzed transcriptomic profiles with RNASeq.

**Main Results:** We found 681 genes that were Mtb-dependent in AMs compared to MDMs and 4538 that were Mtb-dependent in MDMs but not AMs (FDR < 0.05). Using hypergeometric enrichment of DEGs in Broad Hallmark gene sets, we found that Type I and II IFN Response were the only gene sets selectively induced in Mtb-infected AM (FDR <0.05). In contrast, MYC targets, unfolded protein response and MTORC1 signaling, were selectively enriched in MDMs (FDR < 0.05). IFNA1, IFNA8, IFNE, and IFNL1 were specifically and highly upregulated in AMs compared to MDMs at baseline and/or after Mtb infection. IFNA8 modulated Mtb-induced pro-inflammatory cytokines and, compared to other interferons, stimulated unique transcriptomes. Several DNA sensors and Interferon Regulatory Factors had higher expression at baseline and/or after Mtb infection in AMs compared to MDMs.

**Conclusions:** These findings demonstrate that Mtb infection induced unique transcriptional responses in human AMs compared to MDMs, including upregulation of the IFN response pathway and specific DNA sensors.

## Introduction

Alveolar macrophages (AMs) are abundant innate immune cells in the lung and major orchestrators of the inflammation, injury, and repair programs that may dictate the spectrum of tuberculosis (TB) disease (1, 2). Transcriptomic analyses of human macrophages collectively point to baseline RNA profiles that substantially differ between AMs compared to peripheral blood-derived macrophages (3–5). These contrasts in gene expression are likely due to differences in ontogeny as well as environmental exposure. In contrast to blood derived myeloid cells, AMs are constantly exposed to inhaled particles and pathogens which leads to both, activation and downregulation of immune response genes (6). Although prior research documented differences in gene expression profiles between human AMs and MDMs (7, 8), and between tissue macrophages (9), the Mtb-specific AM cellular functions that regulate critical steps in anti-microbial responses have not been identified (5, 10). Understanding AM-specific responses is critical for understanding the pathogenesis of pulmonary pathogens.

Murine models demonstrate that early Mtb infection predominantly targets AM which are more permissive to bacterial growth than bone marrow derived macrophages (BMDM) (11–14). Early innate responses to Mtb in humans are less well-defined and it is not known which Mtb-induced human and murine AM responses are different (15, 16). Previous studies suggest that human AMs inhibit BCG and avirulent H37Ra Mtb growth better than MDMs or monocytes (15, 16). The mechanism underlying this difference is unknown with conflicting data regarding the role of nitric oxide (15, 17). Although these murine and human studies suggest that anti-microbial defenses differ in Mtb-infected AMs and MDMs, the precise mechanisms remain largely unknown.

One of the dominant effector mechanisms that macrophages use in response to pathogenic organisms is the production of Type I Interferon (IFN) (18–20). Blood transcriptional profiles of patients with active TB and those who progress to active TB are enriched for a type I IFN signature (21–23). In human monocyte-derived-macrophages (MDMs), type I IFN inhibits the production of IL1A and IL1B, cytokines that are critical for Mtb host defense (24–26). In addition, an IFNAR1 gene mutation (deletion of the Type I IFN receptor) is associated with increased protection from TB disease (27). Together, these data suggest that Type I IFN signaling is associated with impaired control of Mtb and an increased risk of TB in humans. However, the role of Type I IFN in the lung early responses to Mtb are less clear.

In this study, we tested the hypothesis that Mtb induces a unique and essential anti-microbial pathway in human AMs compared to MDMs by transcriptional profiling of paired human AMs and MDMs from healthy volunteers. We determined that transcriptional profiles were cell-specific and Mtb-dependent in Mtb-infected AMs compared to MDMs. Several Type I IFNs, including IFNA8 and IFNA1, were highly induced in Mtb-infected AMs, but nearly undetectable in MDMs. Together, these data suggest a novel and enhanced Type I IFN response in human AMs that may be a critical component of the early events in TB pathogenesis.

## Methods

### Human Subjects

We recruited 6 healthy, non-smoking, volunteers 18-50 years of age in the Seattle area. Subjects underwent phlebotomy to obtain PBMCs (28–30) and serum as well as fiberoptic bronchoscopy to obtain alveolar macrophages via bronchoalveolar lavage (31). The University of Washington’s human subjects review board approved the study protocols. All participants gave written informed consent.

### Reagents

Roswell Park Memorial Institute (RPMI) 1640 and Dulbecco’s Modified Eagle Medium (DMEM) was purchased from Invitrogen (Carlsbad, CA.). IFNG and IFNB were purchased from Peprotech. IFNA8 purchased from SinoBiological. IFNE was purchased from Bio-Techne.

### Monocyte Derive Macrophages

Human PBMCs for the studies on healthy volunteers that donated blood and AMs were isolated from Vacutainer CPT Cell Preparation Tubes (BD cat no. 362753). PBMCs for the experiments that involved MDMs only were obtained from leukoreduction chambers provided by Bloodwork Northwest (Seattle, WA). PBMCs were isolated by Ficoll gradient separation. Cells were cryopreserved after collection and stored in LN2. Cells were thawed in batch (40E6) and differentiated for 5 days of culture in RPMI/10% FBS with 50ng/mL macrophage-colony stimulating factor (M-CSF) to obtain monocyte-derived macrophages which were purified with magnetic bead separation (CD14 negative isolation protocol, Monocyte Isolation Kit II. Miltenyi Biotec, Auburn, CA). Isolated CD14+ monocyte-derived macrophages (MDMs) were plated at 2E6 cells/well in RPMI/20% autologous serum and maintained in cell culture at 37 °C and 5% CO2 overnight before infection or stimulation.

### Bronchoscopic Collection of Alveolar Macrophages

Fiberoptic bronchoscopy was performed to obtain alveolar macrophages present in the bronchoalveolar lavage from healthy nonsmoking volunteers. None of the recruited volunteers have a history of chronic or acute lung condition. Immediately after the procedure, bronchoalveolar lavage fluid was processed through 70-micron Falcon cell strainers. Cells were pelleted at 300g for 10 minutes at 4C. Cells were washed in 5ml cold Hanks’ Balanced Salt Solution (HBSS)-(without calcium or magnesium) twice and then re-suspended in 5ml RPMI with 10% heat-inactivated FBS, 1% HEPES, 2mM L-glutamine, penicillin 100 U/ml and streptomycin 100 µg/ml. Cell viability was estimated by trypan blue exclusion. Cytospin preparations were stained with a modified Giemsa stain to determine the proportion of macrophages. Alveolar macrophages were plated in RPMI/10% FBS, penicillin and streptomycin at 37 °C, 5% CO2, 95% for 2 hours. After removing non-adherent cells by 6 washings with warm HBSS, adherent cells were then cultured overnight in RPMI/20% autologous serum without antibiotics at 1E6 per well. Alveolar macrophages were treated/infected 24 hours after harvest.

### Cryopreservation of Alveolar Macrophages

For ELISA experiments, cryopreserved AMs were used after assessing various quality control metrics. Standard cryopreservation protocols were used with resuspension of cell pellets of 5E6 AMs/mL in equal volume of RPMI/10%FBS and freeze media (20% DMSO + 80% FBS). For thawing, we slowly warmed the vial in a 37°C waterbath. We slowly pipetted (30seconds) 1 mL of pre-warmed media into the cryovial transferred the contents to a tube of pre-warmed 10mL of RPMI/10% FBS. We washed the cell pellet 2X in fresh media to remove DMSO, counted viable cells with Trypan Blue, and plated AMs at 2E6 cells/mL in RPMI/20% FBS with 1X Pen-Strep (100U/mL) in non-TC treated plates and left cells to rest overnight. The next day we detached any adherent cells with a cell scraper, counted viable AMs with Trypan Blue and plated in a 96-well TC-treated plate at 8.5E4 cells/well in RPMI/10% FBS. We then plate-adhered cells at 37°C/5% CO2 for 2 hours, And then washed with6X HBSS (with Ca/Mg) to remove antibiotics and non-adherent cells. We added RPMI/10% autologous serum and rested them overnight at 37°C/5%CO2.

We compared several quality control measurements in both fresh and cryopreserved AMs. First, we did not observe major differences in morphology or adherence of cryopreserved and freshly harvested AMs in 2 donors (Supplemental Figure 1A). Second, in a comparison of media vs Mtb-induced transcriptional responses, (methods described below) we found high correlation values of gene expression of Mtb stimulated RNA Seq profile of freshly infected AMs when compared to cryopreserved AMs (Supplemental Figure 1B, Pearson R=0.94 – 0.96). Finally, we also found no significant differences in cytokine secretion in Mtb infected fresh AMs vs cryopreserved AMs (Supplemental Figure 1C). Together, these data suggested that cryopreserved AMs have similar functional properties as freshly collected AMs.

### Bacteria preparation and infections for RNA seq

We used Mycobacterium tuberculosis Mtb H37Rv strain (ATCCR 25618TM) for cell infection for RNA sequencing. Mycobacterial stock cultures were thawed and cultured to log-phase in 7H9 media with Middlebrook ADC enrichment (BD), 0.05% Tween 80 (Sigma), and 0.2% Glycerol in a shaking incubator at 37°C and 150RPM in conical tubes with a vented cap. Cultures were washed and reconstituted in Sauton’s media and frozen as stocks. For RNA sequencing, prepared macrophages were infected with freshly thawed Sauton’s washed H37Rv stocks at a multiplicity of infection (MOI) 1:1. After six-hour incubation at 37°C, 5%CO2, cells were lysed and preserved in TRIzol (Invitrogen). and stored at −80C until all infections were completed. TRIzol samples were then thawed, mixed with chloroform (Fisher Scientific), and spun at 12,000g for 15 minutes at 4°C. The upper aqueous layer was mixed with an equal volume of 100% ethanol and processed using miRNeasy Mini Kit (Qiagen) according to the manufacturer’s instructions with the following modifications. DNA digestion was performed with DNAse I (Qiagen) and the final elution was performed once with 30ul RNase-free water to maximize the yield. The average RNA Integrity Number (RIN) as measured by Bioanalyzer was 9.5 and the average yield for AMs (1E6 per well) was 3620 ng. The average yield for MDMs (2E6 per well) was 6720 ng.

### Interferon and LPS treatment of MDMs and AMs

MDMs were treated with recombinant interferons (IFNs): IFNA8 (10ng/mL) (SinoBiological), IFNG (10ng/mL), IFNB (10ng/mL) (Peprotech), and IFNE (100ng/mL) (R&D Systems) for 6 hours before RNA extraction and sequencing. Alternatively, MDMs were treated with IFNs for 24 hours and then infected with H37Rv for an additional 24 hours. Supernatant was then harvested for ELISA. MDMs and AMs were treated with LPS (info) (100ng/mL)

### Cytokine measurements

Macrophages were pretreated with IFNG, IFNB1, IFNA8, or IFNE for 16 hours. and then infected with H37Rv at a MOI of 1:5 for 24 hours. Supernatants were filtered with a 0.22 micron filter to remove samples out of the BSL3. Subsequently, supernatants were evaluated for cytokine concentrations (TNF, IL6 and IL1β) via ELISA Duoset (R&D Systems, Minneapolis, MN). Each sample was assayed in triplicate, and experiments shown were performed at least twice to ensure reproducibility.

### RNA sequency data processing and analysis

For AM vs. MDM and fresh vs cryopreserved AM experiments, libraries were prepared using Clontech SMARTer stranded v2 Pico Mammalian kit (Takara, San Jose, CA) with total RNA (50-100ng) and sequenced with an Illumina NovaSeq 6000at the Genomics Core at the Fred Hutchinson Cancer Research Center in Seattle, WA. For MDM with IFN experiments, libraries were prepared using the NEBNext Ultra II Directional RNA kit (New England BioLabs, Ipswich, MA) and sequenced with an NovaSeq 6000 at the Vanderbilt University Medical Center genomics core (VANTAGE).

Sequence quality was assessed using FastQC (32), and sequencing adapters were removed using AdapterRemoval (33). Reads were aligned to the human genome (GRCh38) using STAR (34). Alignments were assessed with Picard (35) and filtered to paired, non-PCR duplicate, primary alignments with mapping quality > 30 using samtools (36). Reads in genes were counted with Rsubread (37). All samples passed quality filters with median coefficient of variance (CV) coverage < 1, percent pass filter alignment > 85%, and > 8 million reads in genes per sample. There were no apparent PCA outliers or batch effects (Figure 1B).

**Figure 1.**
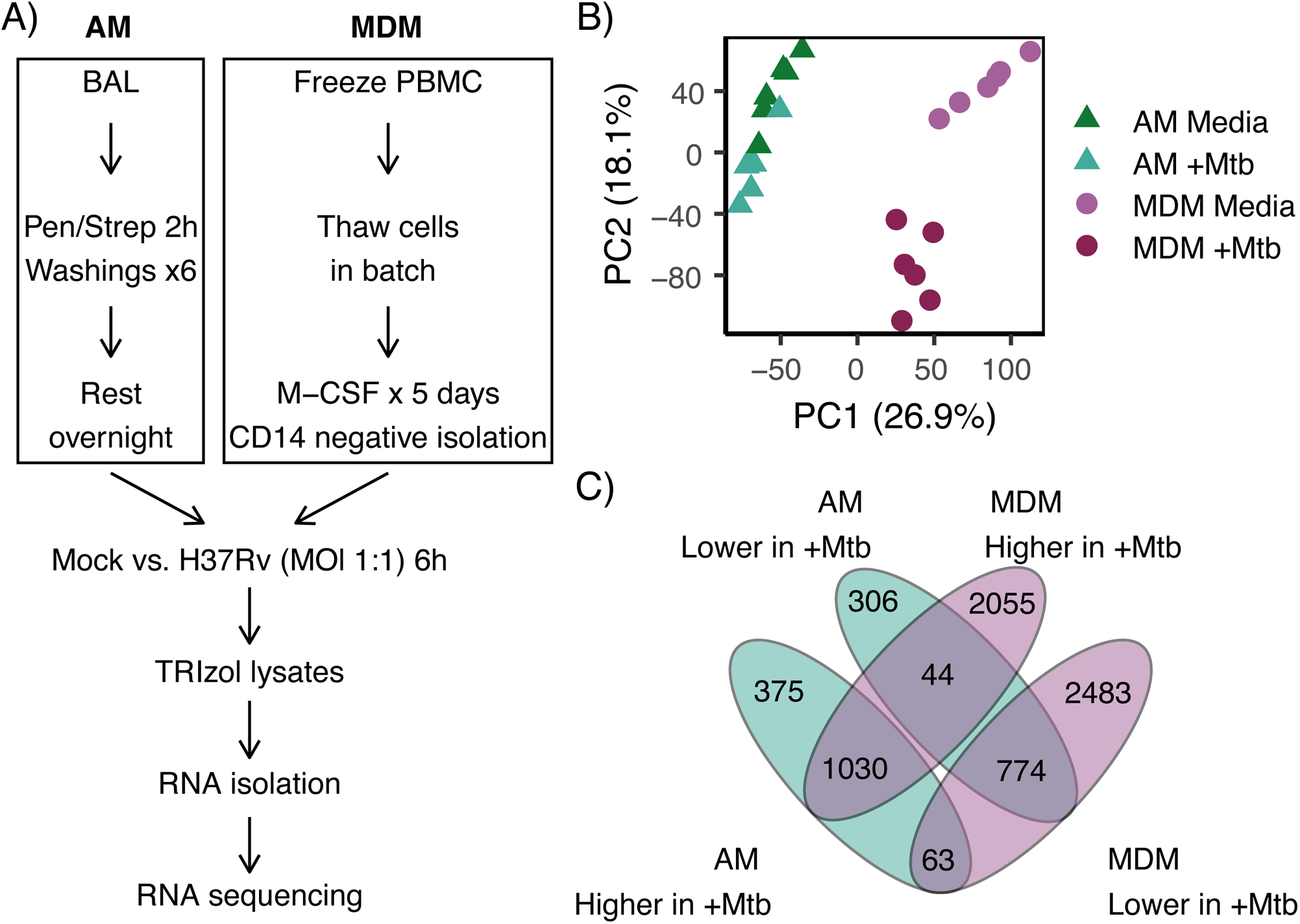
Transcriptional response to Mtb in AM and MDMs. (A) Schematic of study design. Paired AMs and MDMs from 6 healthy individuals were exposed to either media or Mtb strain H37Rv for 6 hours prior to harvesting total RNA for transcriptional profiling with RNAseq (B) Principal component (PC) analysis was performed on log2 normalized expression data. Samples are colored by cell type and Mtb infection. (C) Venn diagram depicting the breakdown of AM and MDM-specific Mtb differentially expressed genes separated by fold change direction.

Data processing and analysis was performed in R v4.2.1 (38) using tidyverse (39), edgeR (40), and limma (41). Protein coding genes were filtered to those with at least 0.1 count per million (CPM) in at least 3 samples, leaving 15,750 (AM vs MDM with Mtb), 15,826 (MDM IFN), and 14,379 (AM cryo vs fresh) genes for analysis. Counts were normalized for RNA composition and voom counts per million (CPM) with quality weights. For the AM and MDM with Mtb experiment, differentially expressed genes (DEGs) were determined using pairwise contrasts between the four-cell type, Mtb treatment groups (AM-media, AM-Mtb, MDM-media, MDM-Mtb) blocked by donor in a linear mixed effects model in kimma (42). DEGs with FDR < 0.05 were defined as those that changed with Mtb-infection in either cell type (AM-media vs AM-Mtb, MDM-media vs MDM-Mtb) and differed between cell types in either infection condition (AM-media vs MDM-media, AM-Mtb vs MDM-Mtb). From these DEGs, six gene lists were curated for AM-specific, MDM-specific, or shared positive or negative responses to Mtb. For the MDM IFN experiment, DEGs were assessed in a linear mixed effects model of IFN blocked by donor. DEGs were defined for each IFN vs media at FDR < 0.05). Gene lists of interest were assessed for enrichment in MSigDB Hallmark gene sets (43) (FDR < 0.05). Gene expression fold change of pairwise contrasts of interest were also assessed for Hallmark in gene set enrichment analysis (GSEA) using fast gene set enrichment analysis (44) and significant pathways were defined (FDR < 0.05). Detailed code is available at https://github.com/hawn-lab/AM_MDM_TB_public

These data have been deposited in NCBI’s Gene Expression Omnibus (GEO) in MIAME-compliant format and are accessible through GEO Series accession number GSE236156.

## Results

### Transcriptional profiles in Mtb-infected human AMs are distinct and enriched for interferons compared to MDMs

To characterize the early human macrophage transcriptional response to Mtb infection, we infected paired human AMs and 5-day M-CSF-derived MDMs from 6 healthy volunteers. At 6 hours after infection, we isolated RNA, and measured genome-wide transcriptional profiles (Figure 1A). Principal components analysis revealed that cell type and infection condition explain variability in the data, mapping strongly to PC1 (26.9% variation explained) and PC2 (18.1%) respectively (Figure 1B). We examined cell-specific and Mtb-dependent differentially expressed genes (DEGs) by using four pairwise contrasts between the conditions (AM-media, AM-Mtb, MDM-media, MDM-Mtb, Supplemental Table I). We found 681 AM-Mtb specific DEGs that were Mtb-dependent in AMs but not MDMs, and 107 that were differentially expressed in both cell types but in different directions, (FDR < 0.05) (Figure 1C, Figure 2, Supplemental Table II). We also identified 4538 MDM-Mtb-specific DEGs that were Mtb dependent in MDMs but not AMs (FDR<0.05) plus the previously described 107 genes with discordant AM and MDM direction (Figure 1C, Figure 2, Supplemental Table II).

**Figure 2.**
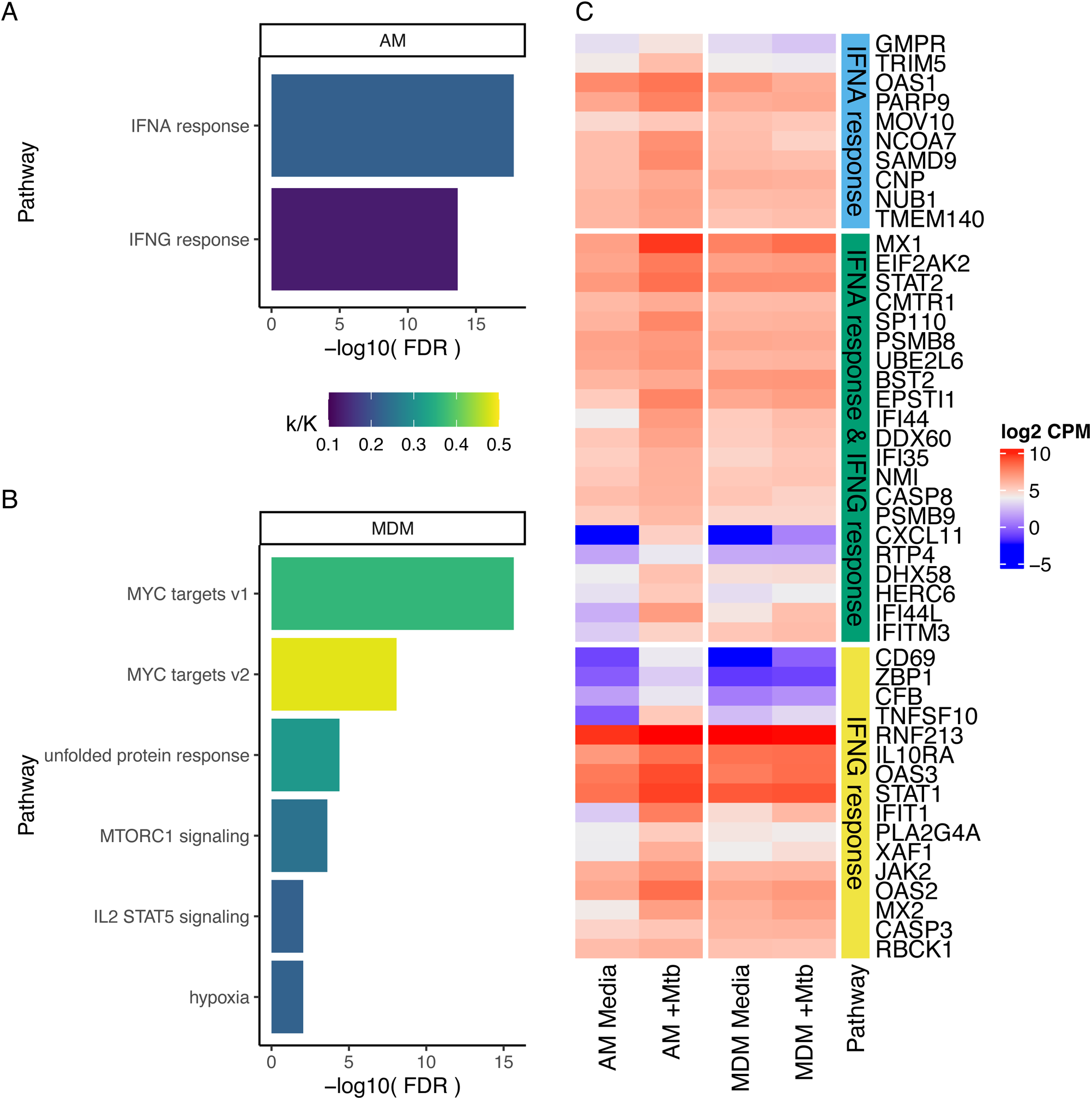
Enrichment of AM-Mtb- and MDM-Mtb-specific DEGs in Hallmark gene sets. Hypergeometric mean enrichment pathway analysis of (A) AM-Mtb and (B) MDM-Mtb specific induced differentially expressed genes (DEGs) using MSigDB Hallmark gene sets at FDR < 0.05. Specific DEGs were defined as significantly upregulated by Mtb-infection in AMs but not MDMs, or vice versa. Color indicates proportion enriched (k/K). (C) Heatmap depicting expression for leading edge genes in MSigDB gene sets significantly enriched by AM-Mtb induced DEGs. Color indicates mean log2 normalized counts per million (CPM) in AM and MDMs with and without Mtb infection. Genes are grouped by presence in one or both of the IFN response gene sets.

We used hypergeometric mean enrichment of MSigDB Hallmark gene sets to characterize AM-Mtb-specific genes with increased expression in response to Mtb (N = 438). The Hallmark Interferon Alpha Response and Interferon Gamma Response were the only gene sets enriched in AMs compared to MDMs (at FDR < 0.1) (Figure 2 and Supplemental Table III). In contrast, hypergeometric enrichment revealed different gene sets, including MYC targets, unfolded protein response, MTORC1 signaling and IL2 STAT5 signaling, highly enriched in MDM-Mtb-specific genes with increased expression in Mtb infected MDMs (FDR < 0.1).

We next used gene set enrichment analysis (GSEA) of Hallmark gene sets to examine the same question across fold changes of the full transcriptome (Supplemental Table IV). In the media condition, there were 5 gene sets that were enriched and more highly expressed in AMs compared to MDMs (Figure 3A, FDR < 0.05). After Mtb infection, there were 6 enriched gene sets with 4 more highly expressed in MDMs including inflammatory response, TNF signaling, allograft rejection, and MTORC1 signaling. In Mtb-infected AMs, there were only 2 gene sets more highly expressed compared to MDMs, and these were the same gene sets discovered in the hypergeometric mean analysis noted above (Interferon Alpha Response and Interferon Gamma Response, FDR < 0.05). Together, these DEG and gene set analyses suggest distinct responses to Mtb in AMs and MDMs with an enrichment of type I and II interferon response pathways in AMs.

**Figure 3.**
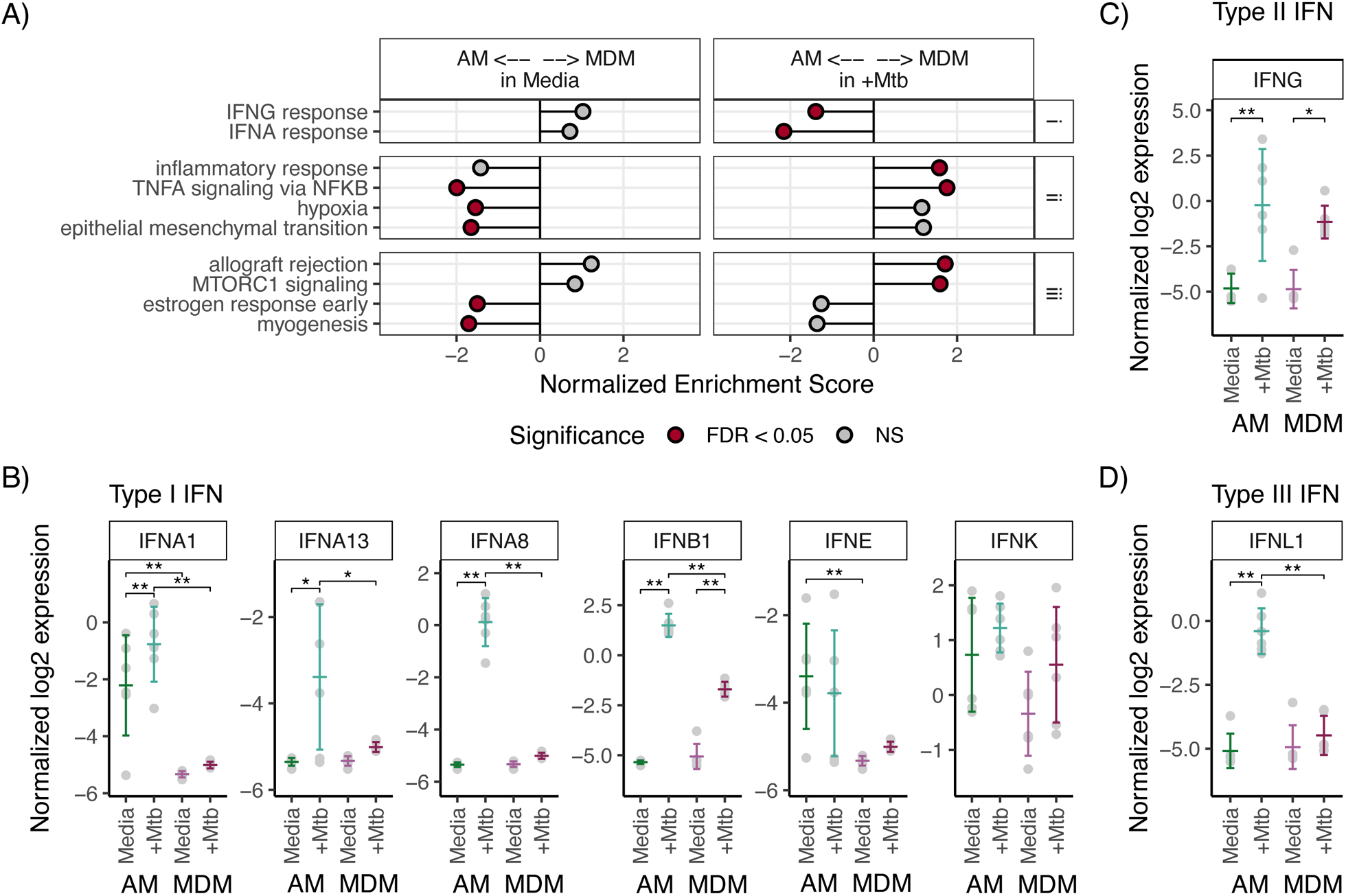
Interferons and interferon response are upregulated in Mtb-infected AMs compared to MDMs. (A) Gene set enrichment analysis (GSEA) with MSigDB Hallmark gene sets comparing AM vs. MDM transcriptional profiles with media or Mtb conditions. Normalized enrichment scores are plotted with color indicating significance (FDR < 0.05). Pathways significant for at least one AM vs MDM contrast as well as at least one Mtb vs media contrast are shown. This resulted in only gene sets with increased expression in response to Mtb infection. Normalized log2 expression of (B) Type I, (C) Type II, and (D) Type III interferons. Bars indicate mean with standard deviation. Pairwise contrast comparisons between AM and MDMs uninfected and Mtb-infected were completed using linear mixed effects modeling. * FDR < 0.05 ** FDR < 0.01.

To identify gene-level differences within the IFN pathways, we next examined expression levels of Type I, II, and III IFNs, their receptors, and other members within the Interferon Alpha Response and Interferon Gamma Response gene sets. We found that IFNB1, three IFNα family members (IFNA1, IFNA8, and IFNA13) and IFNE were upregulated in Mtb-infected AMs compared to MDMs (FDR < 0.05) with minimal to no expression of the remaining 10 IFNAs in either cell type (Figure 3B). There was no difference in expression of IFNG (Type II IFN) between AMs and MDMs though both upregulated IFNG in response to Mtb (Figure 3C). Among the Type III IFNs, IFNL1 was highly upregulated in Mtb-infected AMs compared to MDMs (Figure 3D). Among all the IFN response genes, IFNA8 and IFNL1 were among the top 12% of differentially expressed gene in Ams but not MDMs and, along with IFNB1, were the most differentially expressed IFNS in Mtb-infected AM vs MDM (Supplemental Table I). In summary, these data demonstrate that several interferons are highly enriched in Mtb-infected human AMs compared to MDMs.

### IFNA8 and IFNA1 induction specific to Mtb, but not LPS in AMs

We next examined whether IFNA8 and IFNA1 are induced under other conditions in AMs. We stimulated AMs and MDMs for 6 hours with LPS and measured mRNA levels of IFNA1, IFNA8, IFNB1, IL1B and IL6. As expected LPS induced IL1B and IL6 in both AMs and MDMs. Interestingly, LPS did not induce IFNA1 or IFNA8 in AMs or MDMs. IFNB1 was induced in MDMs but not AMs (Figure 4A). Together, these data suggest that IFNA1 and IFNA8 induction in AMs is dependent on the stimulation condition.

**Figure 4.**
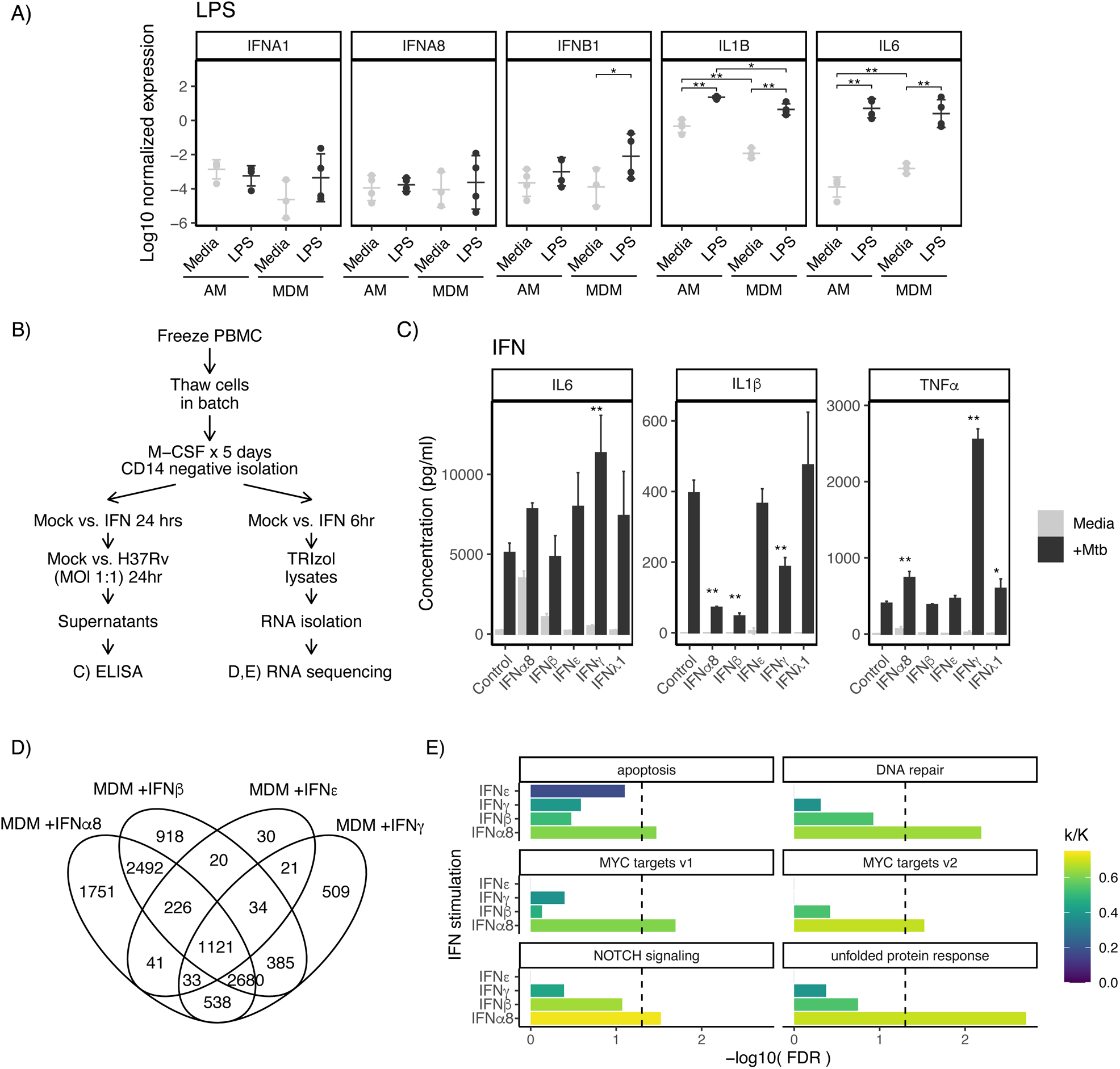
IFNG, IFNB, IFNA8 modulate proinflammatory response in macrophages during in vitro Mtb infection of MDMs. (A) AMs and MDMs stimulated with LPS for 6 hours with IFNA1, IFNA8, IFNB, IL1B and IL6 measured by RT-PCR. Pairwise contrast comparisons between AM and MDMs unstimulated and LPS stimulated were completed using linear mixed effects modeling paired by donor. (**FDR<0.01, *FDR<0.05) (B) Schematic of study design. MDMs from healthy individuals treated with recombinant IFNA8, IFNB, IFNE and IFNG (C) Secreted cytokines: After 24 hours of IFN treatment, cells were infected with H37Rv and TNF, IL6, and IL1B were measured from supernatants harvested overnight. Representative data from two independent experiments are shown. An interaction model of IFN stimulation and Mtb infection was completed using linear mixed effects modeling paired by donor (interaction term **p<0.01, *p<0.05). (D) Cells were treated with IFNs for 6 hours and RNA expression was assessed by RNAseq. Venn diagram depicts IFN-specific and IFN-overlapping DEGs (FDR<0.05). (E) Hypergeometric mean enrichment pathway analysis of IFN DEGs as in (D) against MSigSB Hallmark gene sets. Gene sets specific to a single IFN’s DEGs are shown (FDR < 0.05 for one IFN, FDR > 0.05 for all other IFNs). Color indicates proportion enriched (k/K), and the vertical dashed line indicates FDR = 0.05.

**Figure 5.**
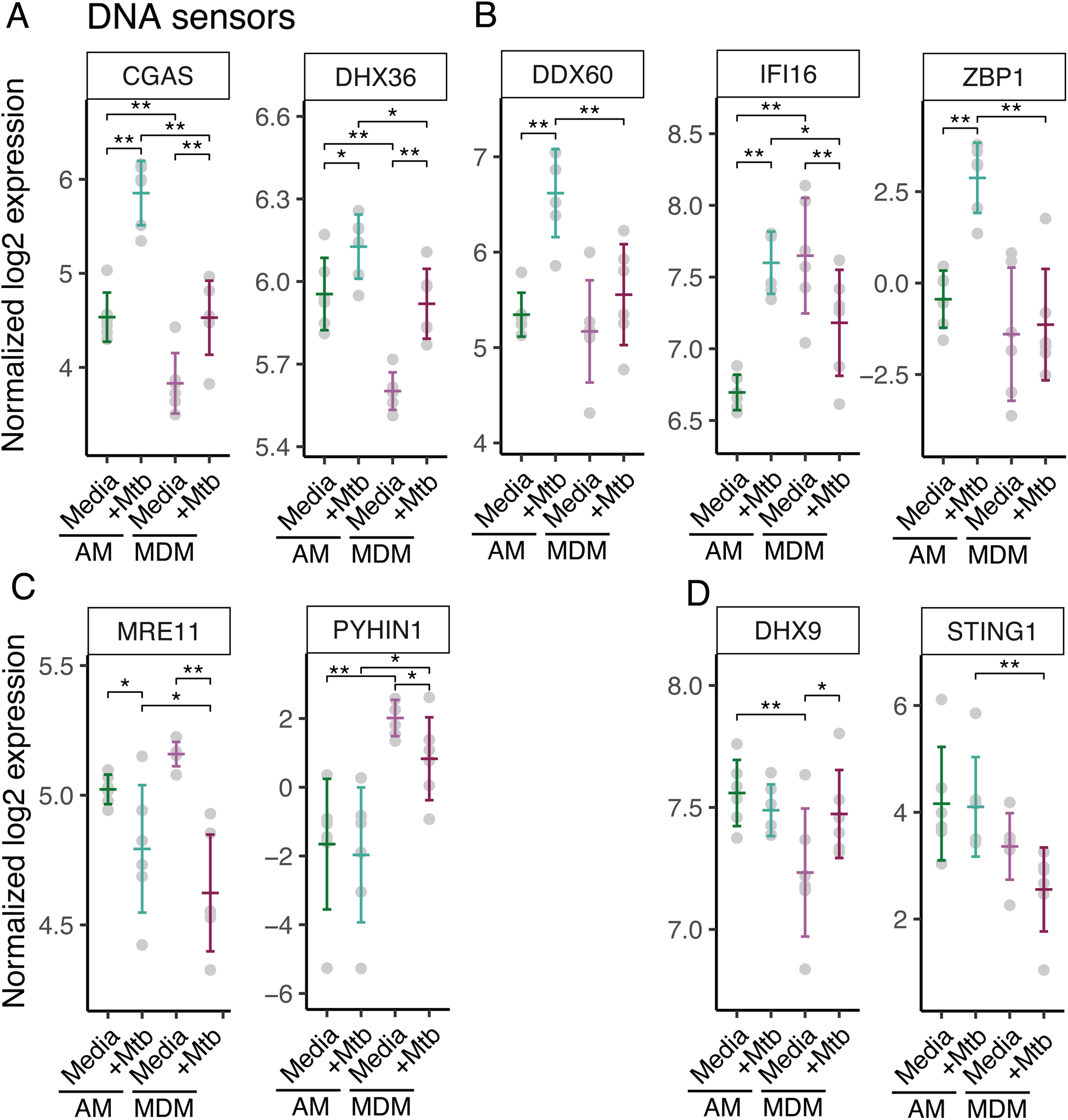
DNA sensor expression at baseline and after Mtb infection in AMs compared to MDMs. Normalized log2 expression of DNA sensor genes from paired AM versus MDM dataset. Bars indicate mean with standard deviation. Pairwise contrast comparisons between uninfected and Mtb-infected AM and MDMs were performed using linear mixed effects modeling. DNA senor genes (A) significantly up-regulated by Mtb infection with higher expression in infected AMs than infected MDMs (cGAS, DHX36), (B) significantly up-regulated by Mtb infection in only AMs (DDX60, IFI16, ZBP1), (C) significantly down-regulated by Mtb infection in AMs and/or MDMs (MRE11, PYHIN1), and (D) not significantly impacted by Mtb infection in AMs (DHX9, STING1). * FDR < 0.05 ** FDR < 0.01.

### IFN-Specific Effects on Mtb-induced cytokine responses in MDMs

To assess whether these IFNs induced different macrophage responses, we measured proinflammatory cytokines from IFN treated MDMs which were infected with Mtb (Figure 4B). In comparison to untreated cells, IFNG induced higher TNF and IL6 levels and decreased IL1B after Mtb infection compared to untreated cells (Figure 4C). IFNB and IFNE induced no differences in secretion of TNF and IL6 compared to untreated cells. In contrast, IFNA8 had decreased secretion of TNF. Similar to IFNG, IFNA8 and IFNB stimulated a decrease in Mtb-induced secretion of IL1B. IFNL1 decreased secretion of TNF and had no difference on IL6 or IL1B. Together these data show that specific IFNs had distinct effects on cytokine secretion in MDMs and specific Type I IFNs had differential effects on cytokine secretion in MDMs.

### IFNA8 Treated MDMs And Mtb-Treated AMs Display Unique Gene Expression Profiles

To discover IFN-specific macrophage signatures, we treated MDMs with IFNG, IFNB1, IFNA8, or IFNE for 6 hours and measured their transcriptional profiles with RNASeq (Supplemental Table V and Figure 4). Each IFN stimulated a mixture of overlapping and distinct transcriptional profiles (Figure 4D). For example, IFNA8 stimulated differential expression of 7131 genes which were also stimulated by at least one other IFN and 1751 genes that were unique to IFNA8 (FDR<0.05). Similarly, IFNB stimulated 918, IFNE stimulated 30, and IFNG stimulated 509 DEGs specific to each IFN. Using hypergeometric mean pathway analysis, we found that the IFNA8 DEGs were enriched in 6 Hallmark gene sets that were not enriched in other IFN DEGs lists (FDR < 0.05), including DNA repair and Unfolded protein response, which were not enriched in any other IFN responses (Supplemental Table VI and Figure 4E). None of the other IFN DEGs yielded IFN-specific enrichments. Together, these data show distinct IFN-specific induced MDM responses including IFNs which bind to the same receptor (IFNA8, IFNE, and IFNB).

### Differential DNA sensor expression in AMs compared to MDMs

To assess possible mechanisms of enhanced IFN signaling in AMs, we next examined whether DNA cytosolic sensors, the major regulators of type I Interferon secretion, are differentially expressed in AMs compared to MDMs. cGAS and DHX36 had increased baseline expression in AMs compared to MDMs. After infection with Mtb, cGAS and DHX36 had increased expression in AMs and MDMs with AMs maintaining higher expression than MDMs (FDR < 0.05) (Figure 6A). Similarly, DDX60, IFI16, and ZBP1 had higher expression in Mtb-infected AMs than MDMs, though these genes were only significantly up-regulated by Mtb infection in AMs (Figure 6B). In contrast, MRE11 and PYHIN1 had lower expression in Mtb-infected AMs and/or MDMs compared to media (Figure 6C). MRE11 was significantly higher in Mtb-infected AMs than MDMs while PYHIN1 was significantly lower in Mtb-infected AMs. Finally, DHX9 and STING1 were not impacted by Mtb infection in AMs but remained moderately more highly expressed in AMs compared to MDMs (Figure 6D).

We also compared the expression of Interferon Regulatory Factors (IRFs) which are transcription factors that mediate Type I IFN signaling. We found that IRF1, 4, 7, and 9 were up-regulated in response to Mtb in AMs and/or MDMs (Figure S2A). IRF1, 7, and 9 had higher expression in Mtb-infected AMs than MDMs while IRF4 was higher in MDMs after infection. IRF3 and 5 were down-regulated by Mtb infection in AMs and/or MDMs with AMs maintaining higher expression after infection (Figure S2B). In contrast, IRF2, 6, and 8 were not significantly impacted by Mtb infection within cell types with IRF6 and 8 maintaining higher expression in AMs than MDMs regardless of infection (Figure S2C). Together, these data demonstrate that expression of several DNA sensors and Type I IFN pathway transcription factors are elevated in AMs compared to MDMs and provide a possible mechanism of how Mtb-induced type I IFN responses are up-regulated in AMs compared to MDMs.

## Discussion

Using transcriptional profiling, we found that interferon response pathways were highly upregulated in Mtb-infected human AMs and a major distinguishing feature compared to MDMs. At the gene level, IFNA8 and IFNL1 were among the top 12% of genes up-regulated by Mtb infection in AMs and along with IFNB1, were the most differentially expressed IFNs in Mtb-infected AM vs MDM. We found higher levels of DNA sensor cGAS at baseline and higher levels after Mtb infection of cGAS, STING1, and IFI16 in AMs compared to MDMs. Furthermore, IFNA8, as well as IFNG and IFNB, but not IFNL1, induced higher levels of TNF, IL6 and lower levels of IL1B in AMs compared to MDMs. Specific IFNs induced differential patterns of gene expression in MDMs. Together, these data suggest a distinct Mtb-induced transcriptional response in human AMs with a new model emphasizing a central role of IFNA8 and IFN response pathways in the early pulmonary response to Mtb infection, which may favor Mtb rather than host control.

Our study supports previous findings that human AM vs MDM transcriptional profiles are different between AMs and MDMs (3–5, 10). We also identify that human AMs mount a stronger type I interferon response at a very early time point (6 hours) after infection when compared to MDMs. Among the type I IFN response, these data indicate that IFNA8 is a major mediator of the early AM response to Mtb infection. In our model, IFNA8 enhanced secretion of TNF and IL6 and decreased IL1b production after Mtb infection. Our data is consistent with prior evidence that different type I IFN activate specific intracellular pathways. Particularly, IFNA8 has been shown to have higher anti-viral potency compared to other Type I IFN subtypes when tested in several human cell lines (45). When compared to IFNA2, IFNA8 did not affect T-cell mobility despite having similar antiviral properties (46). In addition, a single nucleotide polymorphism (SNP) of IFNA8 has been associated with better overall survival of glioma patients (47). That SNP is located in the IFNA8 promoter and has an enhanced activity to bind nuclear proteins including transcription factor Oct1 involved in immune surveillance of glioma progression (48). These examples suggest that subtype-specific intracellular pathways are activated by different type I IFNs. More studies are needed to characterize the role that IFNA8 plays in early Mtb infection in humans and whether it impacts the spectrum of TB outcomes.

Human type I IFNs comprise a family of 17 functional genes and 9 pseudogenes clustered on chromosome 9 that encode 16 proteins: IFNβ, ɛ, −κ, −ω, and 12 subtypes of IFNα (49, 50). Type I IFNs share 35-95% nucleotide sequence identity, but their three-dimensional structures are remarkably similar (51, 52). Despite their homology and common use of a single heterodimeric receptor, the in vitro antiviral and antiproliferative potencies of the subtypes vary (53–56), as do their effects on T-cell and dendritic cell (DC) function (57), and B-cell proliferation (58). Differences among Type I IFN subtypes may reflect (a) their cell type expression (59, 60) (b) ligand affinities for each of the subunits of the IFN alpha receptor (IFNAR1 and IFNAR2) or (c) downstream signaling pathways including interferon regulatory factors (IRF) in particular which may lead to differential transcriptional induction of Type I IFN genes (61). Our data shows that higher IRF3 and IRF7 expression at baseline and after Mtb infection in AMs compared to MDMs may regulate part of the hyper-Type I responsiveness of AMs.

Our study had several limitations. First, we did not include interstitial lung macrophages in our analysis. Interstitial lung macrophages may be a more appropriate control for differentiating lung versus peripheral responses, but they are logistically difficult to obtain from human volunteers. MDMs are a well-studied cell population that serves as a surrogate for peripheral blood myeloid cells. Nonetheless, the phenotype of MDMs vary according to the protocol used for their differentiation (62). Second, a larger sample size might have uncovered additional pathways that differ in AMs and MDMs. However, the high-level cell-specific clustering of the gene expression of the 6 subjects that we analyzed indicates that we obtained robust and highly reproducible results. Third, previous studies show how AM change their phenotype soon after bronchoscopy (5). We choose a time point for our study that balanced the effects of bronchoscopic trauma and resting cells post-bronchoscopy with obtaining a profile as close as possible to the lung environment. Lastly, we do not have evidence that IFNA8 has any connection with TB disease outcomes, which makes this an area for future study.

In summary, this study advances our understanding of the transcriptional heterogeneity between different populations of human macrophages and their response to Mtb infection. We determined that IFNA8 and the type I IFN response in the early stages of Mtb infection are selectively enriched in AMs compared to MDMs and may be crucial in the regulation of subsequent disease outcomes.

## Acknowledgements

We would like to thank the study participants whose invaluable contribution made this project possible. We appreciate the assistance of the skilled bronchoscopy team at Harborview Medical Center in Seattle. We also acknowledge the University of Washington sequencing core for their expertise. We are indebted to the Hawn lab group for insightful discussions and Mr. Darius Jazayeri for editorial support.

## Supplemental Figures and Tables

**Supplemental Figure 1.**
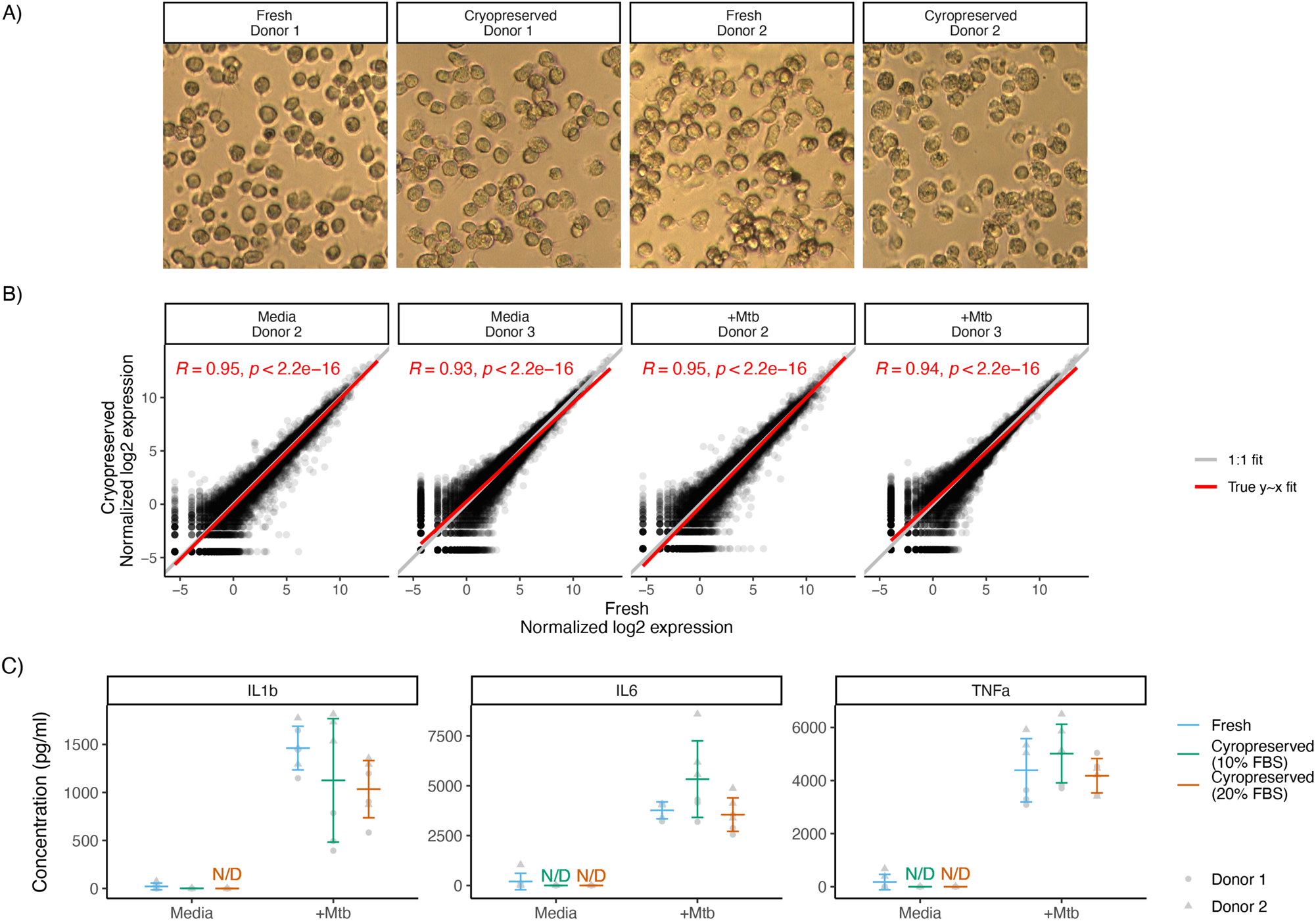
Comparison of cryopreserved vs. frozen human alveolar macrophages. (A) Light microscopy of fresh vs. cryopreserved human alveolar macrophages (AMs) (B) Transcriptional signatures at baseline and after Mtb infection at 6 hours were compared in fresh and cryopreserved AMs. Pearson correlation (R) of gene expression between fresh and cryopreserved AMs with Media and Mtb-infected alveolar macrophages is depicted. The grey fit line indicates a perfect 1:1 fit while the red line is the actual best fit for these data. (R=0.934 – 0.95). (C) Secretion of proinflammatory cytokines from cryopreserved and frozen human AMs. Primary AMs were infected with H37Rv at an MOI of 1:1. IL1B, IL6 and TNFa, were measured from supernatants harvested overnight. N/D: not detected.

**Supplemental Figure 2.**
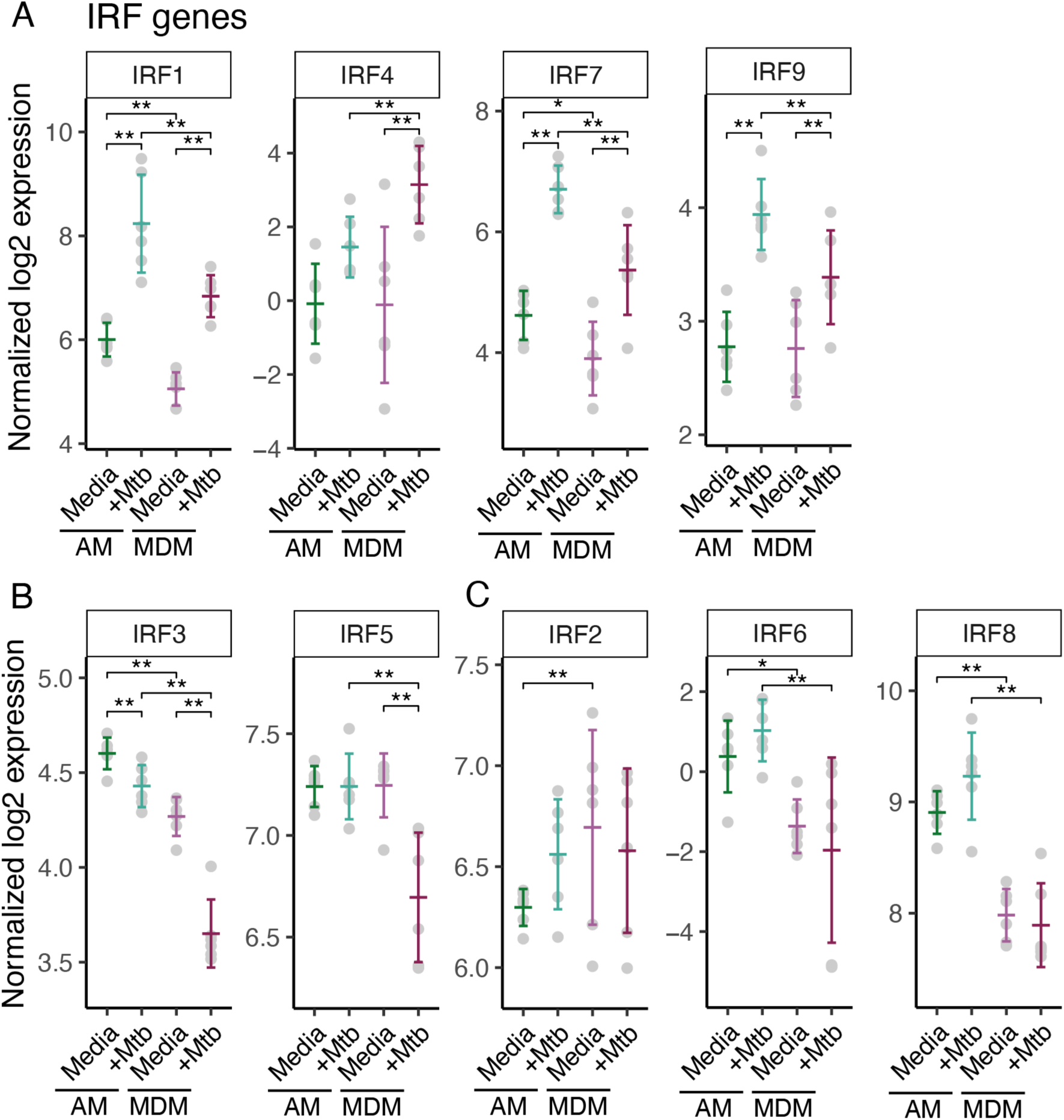
Interferon Regulatory Factor (IRF) gene expression at baseline and after Mtb infection in AMs compared to MDMs. Normalized log2 expression of IRF genes from paired AM versus MDM dataset. Bars indicate mean with standard deviation. Pairwise contrast comparisons between uninfected and Mtb-infected AM and MDMs were performed using linear mixed effects modeling. IRF genes (A) up-regulated by Mtb infection in AMs and/or MDMs (IRF1, 4, 7, 9), (B) down-regulated by Mtb infection in AMs and/or MDMs (IRF3, 5), and (C) not significantly impacted by Mtb infection in either AMs or MDMs (IRF2, 6, 8).* FDR < 0.05 ** FDR < 0.01.

**Table S I. Linear mixed effects models of AM and MDM Mtb infection**. Pairwise contrast models were run for the AM/MDM with Mtb infection experiment (sheets 1-4). Contrasts were calculated as level (contrast_lvl) minus reference (contrast_ref), giving log2 fold changes (estimate) and significance (p-value [pval], corrected p-value [FDR].

**Table S II. AM and MDM-specific Mtb DEGs**. DEGs were defined as genes significant for both 1) AM vs MDM within media and/or Mtb infected samples and 2) Mtb infection within AM and/or MDM (FDR < 0.05). DEG were further parsed by fold change direction and significance for Mtb in AM, MDM, or both (DEG_group). AM-specific DEGs included genes significantly up or down with Mtb infection in only AMs as well as those with opposite fold change directions in AM and MDM. Similarly, MDM-specific DEG were genes significant in only MDM as well as those with opposite fold change directions.

**Table S III. AM and MDM-specific DEG enrichment**. Hypergeometric mean enrichment of AM and MDM-specific Mtb DEGs in MSigDB Hallmark gene sets. AM-specific DEGs included genes significantly up or down with Mtb infection in only AMs as well as those with opposite fold change directions in AM and MDM. Similarly, MDM-specific DEG were genes significant in only MDM as well as those with opposite fold change directions.

**Table S IV. GSEA of AM and MDM with Mtb infection**. Fold change estimates from contrast linear mixed effects models (Table S1) were used in MSigDB Hallmark gene sets with GSEA comparing AM versus MDM with and without Mtb infection. Results are split by sheet for AM vs MDM within media and/or Mtb infected samples and Mtb infection within AM and/or MDM. ES: enrichment score, NES: normalized enrichment score.

**Table S V. Linear mixed effects models of IFN stimulation in MDMs**. Pairwise contrast models were run for MDM treated with IFN stimulation experiment (sheets 1-5). Contrasts were calculated as level (contrast_lvl) minus reference (contrast_ref), giving log2 fold changes (estimate) and significance (p-value [pval], corrected p-value [FDR]).

**Table S VI. IFN-specific DEG enrichment.** Hypergeometric mean enrichment of IFN DEGs in MSigDB Hallmark gene sets. Significant enrichments were defined at FDR < 0.05 for each IFN and IFN-specific genes included those only significant for a single IFN. Only IFNA8 resulted in IFN-specific Hallmark enrichments.

## Notes

Funding: The study was supported by NIH grants K08AI130266, UMN ECRA (M.C.), K24AI137310 (T.R.H.), contract number 75N93019C00070 (T.R.H.), R21AI130798 (S.J.S.) and R21AI156263 (S.J.S.).

### Competing Interest Statement

The authors have declared no competing interest.

